# ASICS: an R package for a whole analysis workflow of 1D ^1^H NMR spectra

**DOI:** 10.1101/407924

**Authors:** Gaëlle Lefort, Laurence Liaubet, Cécile Canlet, Patrick Tardivel, Marie-Christine Pére, Hélène Quesnel, Alain Paris, Nathalie Iannuccelli, Nathalie Vialaneix, Rémi Servien

**Affiliations:** MIAT, Universié de Toulouse, INRA, Castanet Tolosan, France; GenPhySE, Université de Toulouse, INRA, ENVT, Castanet Tolosan, France; Toxalim, Université de Toulouse, INRA, ENVT, INP-Purpan, UPS, 31027 Toulouse, France; Axiom Platform, MetaToul-MetaboHUB, National Infrastructure for Metabolomics and Fluxomics, 31027 Toulouse, France; Institute of Mathematics, University of Wroclaw, Poland; PEGASE, INRA, Agrocampus Ouest, 35590, Saint-Gilles, France; Sorbonne Universités, Muséum National d’Histoire Naturelle, CNRS, UMR7245 MCAM, Paris, France; INTHERES, Université de Toulouse, INRA, ENVT, Toulouse, France

## Abstract

In metabolomics, the detection of new biomarkers from NMR spectra is a promising approach. However, this analysis remains difficult due to the lack of a whole workflow that handles spectra pre-processing, automatic identification and quantification of metabolites and statistical analyses.

We present **ASICS**, an R package that contains a complete workflow to analyse spectra from NMR experiments. It contains an automatic approach to identify and quantify metabolites in a complex mixture spectrum and uses the results of the quantification in untargeted and targeted statistical analyses. **ASICS** was shown to improve the precision of quantification in comparison to existing methods on two independant datasets. In addition, **ASICS** successfully recovered most metabolites that were found important to explain a two level condition describing the samples by a manual and expert analysis based on bucketting. It also found new relevant metabolites involved in metabolic pathways related to risk factors associated with the conditions.

This workflow is available through the R package **ASICS**, available on the Bioconductor platform.

## 1 Introduction

Metabolomics is the comprehensive characterization of the small molecules involved in metabolic chemical reactions. It is a promising approach in systems biology for phenotype characterization or biomarker discovery, and it has been applied to many different fields such as agriculture, biotechnology, microbiology, environment, nutrition or health. Complementary analytical approaches, such as Nuclear Magnetic Resonance (NMR) or High-Resolution Mass Spectrometry, can be used to obtain metabolic profiles. These technologies allow routine detection of hundreds of metabolites in different biological samples (cell cultures, organs, biofluids…). But, due to their high complexity and to the large amount of generated signals, the analysis of such data remains a major challenge for high-throughput metabolomics.

This article focuses on NMR data, that is a promising tool to detect interesting biomarkers. The most common approach to deal with ^1^H NMR spectra is to first divide them into intervals called buckets. The areas under the curve are computed for every bucket and every spectrum and these data are given as inputs to statistical methods to provide a list of buckets of interest (for instance buckets that are significantly different between two conditions). Since buckets are not directly connected to metabolites, this approach requires that ^1^H NMR experts identify the metabolites from the extracted buckets. Not only is this identification step tedious, time consuming, expert dependent and not reproducible but it also leads to a serious loss of information since the identification of metabolites is restricted to the ones that correspond to extracted buckets (Considine *et al*., 2018).

Some methods have thus been developed to automatically identify metabolites from 1H NMR spectra (MetaboHunter (Tulpan *et al*., 2011), MIDTool (Filntisi *et al*., 2017)) and others to automatically quantify the concentration of detected metabolites (Autofit (Weljie *et al*., 2006), **batman** (Hao *et al*., 2012), Bayesil (Ravanbakhsh *et al*., 2015) and **rDolphin** (Cañueto *et al*., 2018)); see Bingol (2018) for a complete review. Recently, Tardivel *et al*. (2017) defined a new statistical method to automatically identify and quantify metabolites that outperforms the other approaches. However, the approach mainly focuses on the quantification step and needed to be embedded in a complete pre-processing and post-processing analysis workflow, available through a simple tool. To our knowledge, such analysis workflows already existed (see a review in Misra (2018)) but they were usually restricted to some steps of the global analysis (post-processing, bucketing or statistical analysis). The only exception seems to be the W4M e-infrastructure (Guitton *et al*. (2017), available through the Galaxy platform^1^), whose automatic identification and quantification step is based on an earlier version of **ASICS** but the environment only allows one-by-one spectrum analysis. Furthermore, none of the existing workflow is as flexible, easily installed and embedded with other tools than an R package can be.

The R package **ASICS** (Automatic Statistical Identification in Complex Spectra) was thus designed to fill this gap. The identification and quantification method is partially based on Tardivel *et al*. (2017) but has been strongly tested, revisited and improved to provide a fine tuning of all the parameters of the original approach.

## 2 Material and methods

**ASICS** is an R package available on Bioconductor^2^ (Gentleman *et al*. (2004)) that combines all the steps of the analysis of ^1^H NMR spectra (library of pure spectra management, preprocessing, quantification, post-quantification statistical analyses). The package also includes functions to directly perform statistical analyses on buckets and diagnosis tools to assess the quality of the quantification. All functionalities of the **ASICS** package are summarized in Figure 1 and described in the next sections.

**Figure 1.**
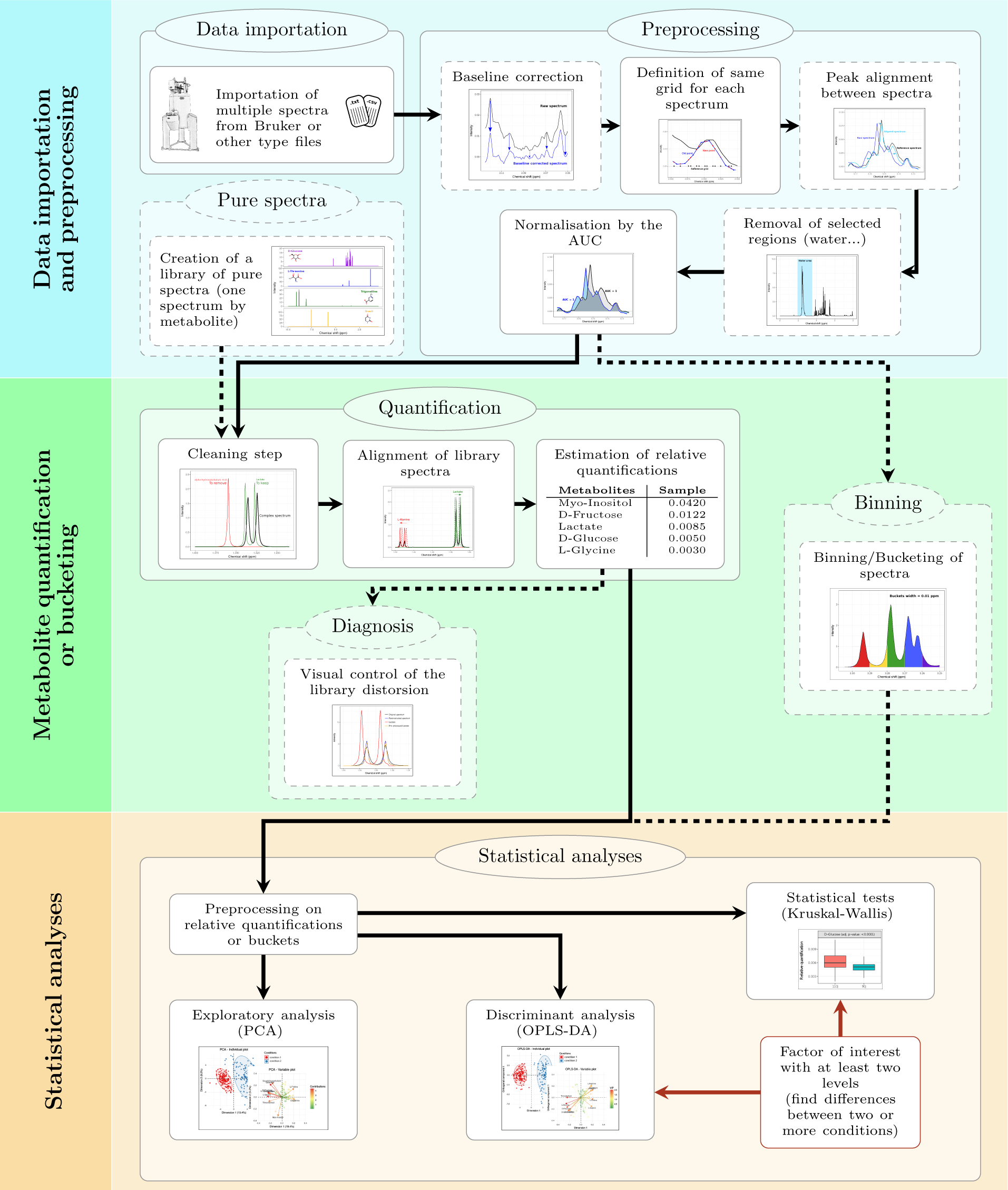
Schematic representation of **ASICS** workflow. Bottom box (with brown background): supplementary data (factor corresponding to experimental conditions for the different spectra) are required for this part of the analysis.

### 2.1 Preprocessing the complex mixture spectrum

After the data are imported from raw 1D Bruker spectral data files or other types of files, several preprocessing steps are recommended in order to remove technical biases.

#### Baseline correction

Most of ^1^H NMR spectra have baseline distortions coming from various sources like instrument instability. These distortions can induce an increase or a decrease in peak intensities and skew the results of quantification. Wang *et al*. (2013) developed a method to estimate the baseline for a spectrum by classifying each point as a signal or a noise point and by using a linear interpolation between noise points to construct the baseline. Then, the baseline is subtracted from its spectrum.

#### Peak alignment

Due to pH or temperature variations between the acquisition of multiple spectra, peak positions of the same metabolite can change between spectra. It is better to align all peaks before analyses, especially if a binning algorithm is used. Vu *et al*. (2011) developed an algorithm, implemented in the R package **speaq**, to carry out this alignment. It is based on continuous wavelet transform to detect peaks and hierarchical clustering to align all spectra on a reference one.

#### Removal of unwanted regions

It is also frequent to exclude a part of the spectra from the analysis. For instance, the part corresponding to water (4.5-5.1 ppm) is of no interest for most biological analyses and thus frequently removed prior to statistical analyses. Urea region (5.5-6.5 ppm) is also frequently excluded in case of urine samples.

#### Normalisation

A normalisation is mandatory before any analysis to make samples comparable. It allows to minimise systematic variations due to differences in sample dilutions. One of the most used methods is the normalisation to a constant sum (Craig *et al*. (2006)). As a result, the total spectral intensity is the same for each spectrum.

In **ASICS**, all preprocessing steps are available and the normalisation is the only mandatory one (it is systematically performed when the data are loaded). In the two following steps of the quantification method (preprocessing of the reference library, described in Section 2.2, and quantification itself, described in Section 2.3), all complex mixture spectra are processed individually and independently from each other. The method is thus described for only one complex mixture spectrum (and repeated similarly for all the others).

### 2.2 Preprocessing the reference library

A library of pure metabolite spectra is used as a reference to identify and quantify metabolite concentrations in the (complex mixture) spectra of interest. This library is a set of spectra of pure compounds, that have been acquired independently from samples. Such a reference library is available in **ASICS**. This library is composed of 190 spectra for which the noise has already been removed (Supplementary Data 1). The spectra acquisition procedure is detailed in Tardivel *et al*. (2017). In addition, **ASICS** provides functions to add or remove some spectra from the available reference library or to use another (user provided) reference library.

In addition to removing noise of each library spectrum, preprocessing steps are needed to clean and adapt the library to each spectrum of interest.

#### Noise thresholding

As this is the case for each ^1^H NMR spectrum, all spectra in library contain noise. All values below a certain threshold, *s*_*l*_, (that can be defined by the user; default value is *s*_*l*_ = 1), are considered as noise and set to 0. This allows to select peak positions, a step that is critical for the next selection stage.

#### First selection step

A metabolite can not belong to the complex mixture if at least one peak of its spectrum does not appear in the complex mixture spectrum peaks. Using this simple property, a first selection step is performed. All spectra in the reference library for which the peaks are not included in the peaks of the complex mixture spectrum are removed. This step results in a reference library of *p* pre-selected reference spectra that are used in the model described in Section 2.3. As technical biases can yield to chemical shifts, a reference spectrum is selected if all its peaks are present in the complex mixture spectrum with an allowed shift of *M* ppm between the two spectra. In addition, as complex mixture spectra are noisy, peaks under a threshold *s*_*m*_ are ignored for this identification step. By default, the maximum allowed shift is *M* = 0.02 ppm and the threshold is *s*_*m*_ = 0.02. However, both values can be changed by the user, depending on his spectrometer and experimental conditions.

#### Translation and distortion

The alignment algorithm described in Section 2.1 can not be used to align reference spectra with the complex mixture spectrum. The reason is that this method is not adapted to spectra with a low number of peaks as those of the pure metabolite contained in the reference library. A two step procedure is used instead (Figure 2):

**Figure 2.**
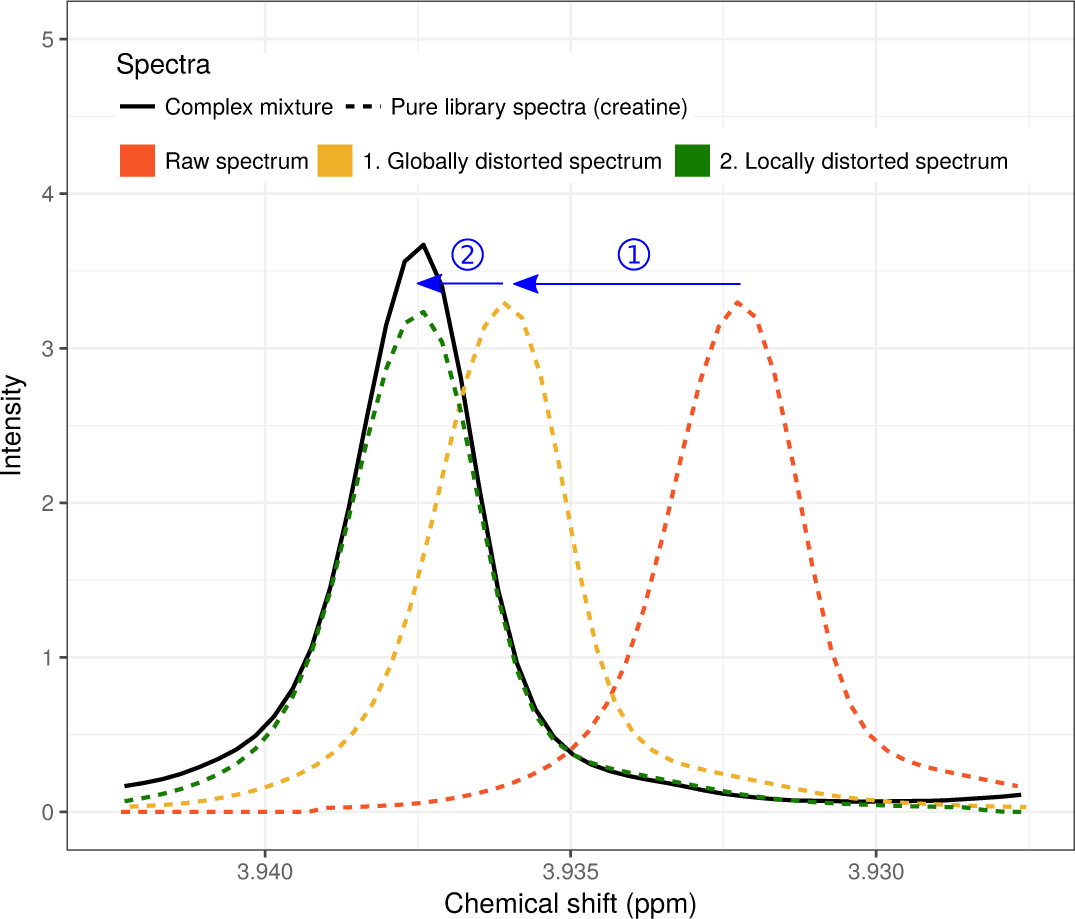
Two steps distortion procedure for the main peak of the creatine. 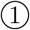 Global translation of the creatine spectrum. 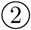 Localized distortion of one of the creatine peak.

1. First, reference library spectra are aligned with the complex mixture spectrum of interest by maximizing the Fast Fourier Transform cross-correlation *(Wong et al. (2005)).* The algorithm that finds the best shift (with a maximum allowed shift equal to *M*) is taken from the R package **speaq** (Vu *et al*. (2011)).
2. Second, every peak of each library spectrum taken individually is aligned by minimizing residuals of the linear regression between the spectrum of interest and the reference library spectrum. To perform local distortions of the chemical shift grid for each peak, **ASICS** uses the function *ϕ*(*x*) = *ax*(1 - *x*) + *x*, where *x* ∈ [0, 1], corresponds to the rescaled initial grid, *ϕ*(*x*) *∈* [0, 1] to the newly scaled grid and 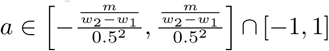 is a coefficient of distortion. The definition domain of *a* is controlled by *m*, the maximum allowed shift (with 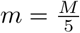) and by (*w*_1_*, w*_2_) that are the lower and upper bounds of the initial grid, respectively. Different distortions are tested for each peak and only the one that minimizes the residuals is kept to build the final reference library used in the quantification algorithm.

### 2.3 Metabolite quantification

Using the preprocessed complex mixture spectrum and the preprocessed spectra of the reference library, the metabolite identification and quantification in the complex mixture spectrum is performed as described in Tardivel *et al*. (2017). The complex mixture spectrum is defined as a linear combination of the library reference spectra:

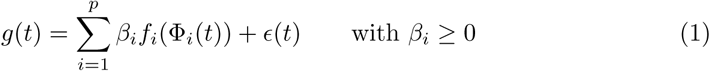

where *g* corresponds to the complex mixture spectrum, *f*_*i*_ ∘ Φ_*i*_ to the *p* pre-selected pre-processed spectra of the reference library, *β* = (*β*_1_, …, *β_p_*) to the coefficients associated with these spectra (or, equivalently, with the corresponding metabolites) and *ϵ* to the noise. The noise is structured so as to take into account both an additive noise, *ϵ*_2_, and a multiplicative noise, 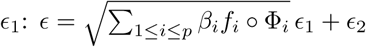.

A variable selection procedure is implemented to obtain a sparse *β* by controlling the Family Wise Error Rate (FWER) with a risk *α*. Usually, the threshold for rejecting ℋ _0_: *β_i_* = 0 is the same for every *i*. Here, we used the procedure described in Tardivel (2017) that allows to define metabolite dependent thresholds in order to maximize the test power. More precisely, the custom thresholds *b*_*i*_ are computed to minimize the volume of the acceptance region, namely 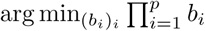 subject to 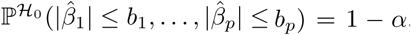, where 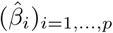 are MLE estimates of Equation (1). The solution of this optimization problem is obtained by simulating a large number of realizations of the random variable *Z* ∼ 𝒩 (0, Σ), where Σ is the estimated variance of the estimates 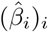 so as to have 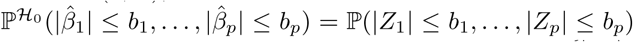, and the thresholds (*b*_*i*_) are obtained as the 1 − *α* quantile of the random variable {|*Z*_1_|, …, |*Z*_*p*_|}, that allows to control the FWER.

Once the metabolites selected, the quantifications (*β*_*i*_)_*i*_ for those selected metabolites are re-estimated by restricting Equation (1) to this subset in order to limit estimation bias. Finally, the relative quantifications are obtained by dividing 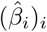 by the respective number of protons of each selected metabolite.

In **ASICS**, pure library preprocessing and quantification are implemented in a unique function that can be run at once for several spectra with a parallel computing backend.

### 2.4 Post-quantification statistical analyses

On quantified metabolites (or on a subset of metabolites that are sufficiently frequently observed in the whole set of complex mixture spectra), the following analyses can be performed, using the spectra alone or a two level factor of interest that describes the samples:

#### Quantification assessment

To assess the quality of **ASICS** quantification, a plot with the original complex mixture spectrum, *g*(*t*), and the reconstructed spectrum, 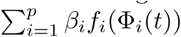, can be obtained for a given sample (Supplementary Figure S1). In addition, one reference spectrum for a given metabolite, and its distorted spectrum, can be superimposed to this plot in order to assess the quality of the metabolite selection for metabolites of interest.

#### Exploratory analysis

To explore results and detect outliers or batch effects, Principal Component Analysis (PCA) can be performed. Individual and variable plots are available to ease the visualisation and interpretation of PCA results (Supplementary Figure S2).

#### Discriminant analysis

When the question of interest is to find which metabolites are different between two experimental conditions describing the samples, Orthogonal Projections to Latent Structures Discriminant Analysis (OPLS-DA, Trygg and Wold (2002)) can be performed with a dedicated function based on the implementation available in the **ropls** package (Thévenot *et al*., 2015). To avoid overfitting, a 10-fold cross-validation procedure is implemented. Prediction error and variable importance in projection (VIP) are computed on ten OPLS-DA and averaged over the ten folds. Moreover, a stability index is created to find influential metabolites (*i.e.*, metabolites with a VIP superior to a predefined threshold) common to most folds. As for PCA, individual and variable plots are available (Supplementary Figure S3).

#### Statistical tests

To answer the same type of questions, statistical tests have also been implemented. Since relative quantifications are usually non normally distributed, Kruskal-Wallis tests are used to find differences between the two (or more) groups, in combination with a correction for multiple testing, as available in the R function p.adjust. Boxplots showing the differences in metabolite quantification between the conditions can be displayed (Supplementary Figure S4).

## 3 Case studies

### 3.1 Plasma metabolome at the end of gestation in piglets

Genetic selection performed during the last decades has been associated with an increase in perinatal mortality in domestic pig, *Sus scrofa* (Canario *et al*., 2006, 2007). One main factor related to neonate survival is the maturation of fetal tissues and organs in late gestation (Gondret *et al*., 2018; Voillet *et al*., 2014, 2018; Yao *et al*., 2017). In order to explore the development of the metabolic status in late gestation, an experiment was performed on pig fetuses. Metabolomic data were acquired on plasma samples collected on *n* = 155 Large White (LW) fetuses at 90 days of gestation and on *n* = 128 fetuses at 110 days of gestation (birth is around 114 days; ANR PORCINET project). All ^1^H NMR spectra were phased and baseline corrected. Glucose, fructose, and lactate were directly quantified by standard methods (they have been chosen as indicators of carbohydrate metabolism). More details about the experimental design and data acquisition can be found in Supplementary Section S2.

Similar analyses were performed on buckets and on relative quantifications computed with **ASICS** to assess the performance of the method. The aim of these analyses was to find differentially concentrated metabolites between two groups: fetuses at 90 and 110 days of gestation. Lists of metabolites obtained with both approaches were compared as well as the direction of change between groups, based on two OPLS-DA, one on buckets and the other on quantifications obtained with **ASICS**. VIP thresholds for both OPLS-DA were set to 1.

Metabolites that were quantified were used to make a quantitative assessment of **ASICS** by comparing the obtained (estimated) quantifications with the dosages. Pearson correlation between quantifications and dosages were computed for every metabolite directly measured by dosage. These correlations were also compared with the correlations obtained for other quantification methods: Autofit, **batman**, Bayesil and **rDolphin**. Contrary to **ASICS**, these methods were too slow or not automated to allow the quantification for the 283 spectra. Therefore, quantifications were performed on a subsample of the original dataset that corresponded to the deciles of the lactate, fructose and glucose dosage to ensure representativity (32 spectra). Computational times were also recorded. For **ASICS** quantifications, water and urea regions were excluded and the maximum shift, *M*, was set to 0.01. In order to perform all quantifications with **batman** in a reasonable time, its library was reduced to the 160 common metabolites between **batman** and **ASICS** reference libraries and the number of iterations was set to 10,000.

### 3.2 Urinary metabolome of Type 2 diabetes mellitus

In order to test our method on data acquired with another spectrometer than the one on which the pure metabolite library included in **ASICS** has been obtained, we used the public datasets from Salek *et al*. (2007). The experiment has been designed to improve the understanding of early stage of type 2 diabetes mellitus (T2DM) development. 1H NMR human metabolome was obtained from 84 healthy volunteers and 50 T2DM patients. Raw 1D Bruker spectral data files were found in the MetaboLights database (Haug *et al*. (2013); study MTBLS1). In the original study, spectra were normalized by the area under the curve after excluding water (4.24–5.04 ppm), urea (5.04–6.00 ppm) and glucose (3.19–3.99 ppm, 5.21–5.27 ppm) regions. Finally, a bucketing was performed with a 0.04-ppm width. The original study used a combination of PLS-DA and statistical tests (*t*-test, *F* -test, Kruskal-Wallis test and Kolmogorov-Smirnov test) on buckets (with a manual expert identification) to find differences between the healthy and ill individuals. This dataset allowed us to test the performance of **ASICS** on a different fluid (urine) in a different species (human).

Contrary to Salek *et al*. (2007), we kept glucose region for a quantification with **ASICS** because the glucose spectrum was available in the library. However, regions of water and urea were excluded. The other parameters of the different methods were set to their default values except for **ASICS** threshold that was set to *s*_*m*_ = 0.05, because we had observed that this dataset was noisier than the previous one. In addition, to control differences that could originate from the differential analysis method itself, we performed the comparison between the buckets and the ASICS quantifications approaches with OPLS-DA, as for the study about perinatal survival (VIP thresholds set to 1.2).

## 4 Results and discussion

### 4.1 Comparison with biochemical dosages on piglets

Correlations between quantifications and biochemical dosages of the three metabolites were performed on the 32 selected spectra. We were not able to obtain quantifications with Bayesil because no chemical shift reference (TSP) has been added during spectrum acquisition. Bayesil handles spectrum from raw NMR induction-decay signal and so it requires that spectra are collected with TSP added to the sample (Beirnaert *et al*., 2018; Ravanbakhsh *et al*., 2015), TSP is sometimes used as an internal reference in samples for NMR. This procedure is not advised for plasma metabolome, and thus not routinely applied, since TSP binds to plasma proteins (Beckonert *et al*., 2007).

Table 1 provides the correlations between the quantified target metabolites and their corresponding dosages for the different quantification methods. In addition, the table includes the correlation between one bucket of the target metabolite and the corresponding dosage as a reference value. These results show that **ASICS** outperforms Autofit, **batman** and **rDolphin** for the three metabolites and provides quantification whose correlations are identical to the ones obtained with a direct comparison to the buckets. Results obtained with **batman** and the library with 160 metabolites are consistent with findings of other studies: the method is not suited for untargeted approaches (Beirnaert *et al*., 2018; Tardivel *et al*., 2017). If the quantification with **batman** is performed including only the three targeted metabolites in the reference library, correlations become similar to the ones obtained by the other methods, but are still lower than those obtained by **ASICS** with no prior selection of the reference library.

**Table 1.**
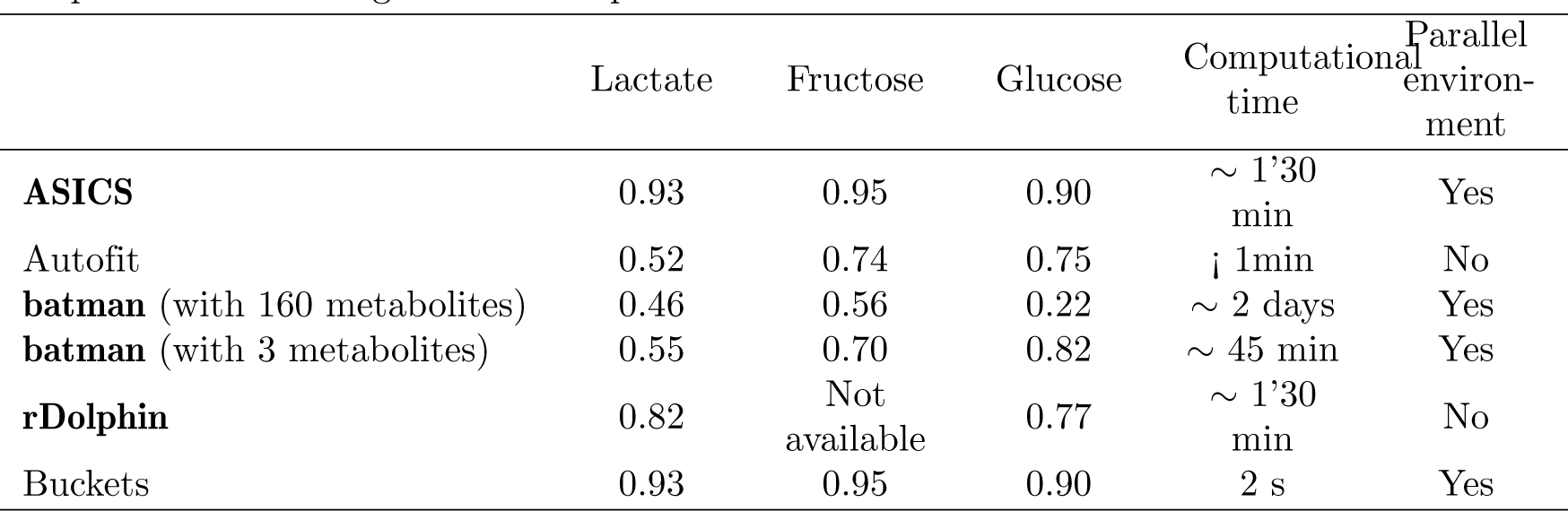
Correlation between biochemical dosages of three metabolites and relative quantifications obtained with four methods and the buckets. Bucket for lactate: 1.335; bucket for fructose: 3.995; bucket for glucose: 5.235. Computational time is given for one spectrum.

On a practical point of view, **ASICS** has other interesting features: first, it provides an easy way to handle (complement, replace, manipulate) the reference library whereas **batman** and **rDolphin** need that information on each multiplet (chemical shift position, multiplicity…) is specified. A biochemical expertise is thus required for the modification of the reference library in these packages. Autofit is a commercial software that requires the acquisition of a license, which strongly limits its use. Finally, the reference library cannot be modified in Bayesil and this method is only available through a web interface that makes automation of several spectra processing impossible.

In terms of computational times, the preprocessing of the library and the metabolite quantification with **ASICS** takes about 1’30 min per spectrum and can be launched at once in parallel. A parallel environment is also available for **batman** but the quantification of a single spectra takes approximately 2 days. Computational time needed by **rDolphin** is approximately the same than for **ASICS** but parallel implementation is not proposed in the package. Only Autofit has a lower computation time than **ASICS** (less than one minute) but spectra can only be quantified sequentially (no parallel environment).

A table summarizing capabilities of each method is available in Supplementary Table S3.

### 4.2 Differences between gestational ages of fetuses

For the study about fetuses in late gestation, two outliers were detected on the bucket dataset in a preliminary study (Supplementary Figure S5) and were removed from the analysis (Supplementary Figures S6 and S7).

OPLS-DA was performed on quantified metabolites and on buckets. Both showed the same predicting power: all samples were perfectly separated according to their stages of gestation. For the bucket analysis, VIP values identified 268 buckets on 781 that were found influential to separate the two groups. Based on this list, a manual identification performed by an NMR expert hightlighted 21 metabolites.

The same analysis was performed on the **ASICS** quantifications and allowed to obtain 22 metabolites. The results obtained by **ASICS** and buckets analysis are detailed in Supplementary Table S4. Nine metabolites were found common to both analyses (Supplementary Figure S8): lactate, creatinine, fructose, glucose, threonine, valine, alanine, proline and leucine. For the metabolites which were not identified by both approaches, we observed five cases:

- metabolites only identified on buckets because the pure spectra was not present in the **ASICS** reference library: the 3-methyl-2-oxovaleric acid and the lipids;
- metabolites that were not found by **ASICS** because differences between the two groups are significative with Kruskal-Wallis tests but not found large enough to be selected by the OPLS-DA: the betaine and the glutamic acid;
- metabolites with low intensity peaks because **ASICS** was not able to identify and quantify smaller quantities: citrate, tyrosine, lysine, creatine and isoleucine;
- metabolites that were found by **ASICS** and not by the bucket analysis but for which all peaks corresponded to buckets that were found influential in the bucket analysis: the glycine and the guanidinoacetic acid. For this case, it is very likely that the non identification of these metabolites comes from an expertise bias (peaks are confused with glucose and fructose thus the expert does not identify it);
- metabolites for which no clear conclusion could be driven on their presence without expert knowledge in NMR or biology or the help of other technologies like 2D NMR spectrometry. For **ASICS** analysis these metabolites correspond to metabolites whose spectra have peaks only in the region with a high density of peaks (3.5 to 4.2 ppm; threonic acid, xylitol, sorbitol, galactitol, glucolic acid and arabitol), with a low concentration (N-acetylglycine, acetamidomethylcysteine, arginine and isovaleric acid) or with peaks confused with glucose peaks (glucose-6-phosphate).

The metabolites found by **ASICS** are consistent with known biological processes of late gestation in pig, especially with the fetal two-fold increase of weight during the last three weeks. It is expected to find up-regulation of the protein synthesis in late gestation, which is illustrated by the increase of amino acid abundances (alanine, proline, threonine, arginine, leucine, valine) just before birth. Also, functional analysis performed with IPA (see Supplementary Figure S9) highlighted 13 metabolites (among the 22 identified by **ASICS**) involved in common metabolic pathways directly related to late stage gestation (survival or organism, metabolism of protein, conversion of lipid). Among these metabolites, 6 (guanidinoacetic acid, sorbitol, glucose-6-phosphate, glycine, gluconic acid and arginine) were identified only by **ASICS** and not with the bucket approach. In this study, the only weakness of **ASICS** is thus a tendency to miss low concentrated metabolites, especially if those have peaks only in the region with a high density of peaks.

### 4.3 Differences for T2DM patients

Results for the T2DM study are provided in Supplementary Table S4 and in Supplementary Figure S10. The same conclusions than in Section 4.2 can be driven: some metabolites were extracted by both analyses (creatinine, betaine, hippuric acid, guanidinoacetic acid, alanine, glucose, indoxylsulfate, acetoacetate and trigonelline), some did not have a pure spectra available in the library (phenylacetylglycine and 2PY) and the **ASICS** algorithm had difficulties to identify metabolites with low concentrations (3-hydroxybutyrate, isoleucine, 2-oxoisovalerate, fumaric acid and butyrate) or with only one proton (allantoin).

In addition, results were compared with those previously obtained by Salek *et al.* (2007) with the same NMR data and with those of an independant experiment realized on urine samples (among other samples) from T2DM patients with another non targeted metabolomic technology Yousri *et al*. (2015) (results also given in Supplementary Table S4). Those comparisons highlighted the relevance of **ASICS** quantification that showed results consistent with previous studies and prior knowledge on Type 2 diabete: some of the metabolites were extracted by **ASICS** and by bucket quantification, like alanine or acetoacetate (Supplementary Table S4), and have also been identified in Salek *et al*. (2007). We were also able to extract other metabolites, like the glucose (D-Glucose), the guanidinoacetic acid or the glycerol, that were not previously described because the glucose region was excluded from the study in Salek *et al*. (2007). The glycerol was identified both by **ASICS** and by Salek *et al*. (2007) in experiments on rats and mice (the glucose region was only excluded in the human dataset and not in the rat and mouse datasets). In all experiments, the glycerol increased in diabetics, which might reflect changes in fatty acids metabolism. With **ASICS**, the creatinine and its precursor, the guanidinoacetic acid (both also found with buckets), were directly quantified in urine and only the creatinine was previously described in (Salek *et al*., 2007; Yousri *et al*., 2015) as down regulated in T2DM. Both these metabolites reflect possible impairment of the renal function in diabetics.

In addition, three metabolites (acetoacetate, acetone and 3-hydroxybutyrate) reflected the presence of ketone bodies in urine when complications for diabete are likely to occur (Misra and Oliver, 2015). 3-hydroxybutyrate and acetoacetate are detected by Salek *et al*. (2007) and Yousri *et al*. (2015) together with buckets and **ASICS**. Acetone is only identified as discriminant by **ASICS** allowing the possibility to reflect the risk of acidocetose in diabetics. Another metabolite rarely identified in T2DM, arabitol (L-Arabitol), was quantified as decreasing only with **ASICS** and firstly described by Yousri *et al*. (2015) in urine of patients. Together with glucose-6-phosphate, also only identified with **ASICS**, these metabolites reflect the pentose pathway activity in diabetics. Metabolites associated to this pathway were also previously identified in urine as strongly associated with T2DM development in a diabetic rat model (Sun *et al*., 2014). Finally, only ASICS allowed the identification of GABA (*Γ*-aminobutyric acid), a neuromediator recently identified to be increased in T2DM and related to a lower cognitive functioning observed in some diabetic patients (Van Bussel *et al*., 2016).

In conclusion, not only was **ASICS** able to automatically recover the main findings of the bucket and expert analysis, it was also able to extract a number of metabolites that are relevant and confirmed by other independent studies but not found by the bucket and expert analysis (glycerol, guanidinoacetic acid, acetone, arabitol, glucose-6-phosphate and GABA). This untargeted approach allowed to highlight several metabolic pathways linked to Type 2 Diabete Mellitus, as illustrated in Supplementary Figure S11.

## 5 Conclusion

This article presents an R package, **ASICS**, integrating a complete analysis workflow of 1H NMR spectra. This pipeline integrates an automatic metabolite identification and quantification method based on a reference library of pure metabolite spectra. **ASICS** showed better quantification results than existing methods and allowed to perform a complete study on several hundreds spectra in only a few hours. Its use on two real world datasets exhibited similar results than the standard analysis on buckets followed by expert manual identification but also allowed to provide new information. For both studies, new metabolites, not extracted by expert identification, were found by **ASICS**, some of them confirmed by previous and independent studies. Obviously, as is the case for other omics data, in coming to a conclusion on whether the metabolites were really present in samples, a validation would be necessary. This validation could be performed with other technologies, like 2D NMR spectrometry, or specific biochemical dosages but is out of the scope of the present article.

Finally, **ASICS** still has some limitations: the algorithm had difficulties to identify metabolites with low concentrations or with their peaks all located in a region with a high density of peaks. Future work will tackle this aspect, by trying to couple the information from the whole set of spectra to improve the individual quantification.

## Funding

This project received financial support from French National Agency of Research (PORCINET grant ANR-09-GENM005). The PhD fellowship of Gaelle Lefort is supported by the Digital Agriculture Convergence Lab (#DigitAg, http://www.hdigitag.fr/), by the INRA Mathematics and Computer Science Division, by the INRA Animal Genetics Division and by the INRA Animal Health Divison.

## Footnotes

1 https://usegalaxy.org/

2 http://bioconductor.org/packages/ASICS/

